# Development of an N-Cadherin Biofunctionalized Hydrogel to Support the Formation of Synaptically Connected Neural Networks

**DOI:** 10.1101/729079

**Authors:** Brian J. O’Grady, Kylie M. Balotin, Allison M. Bosworth, P. Mason McClatchey, Robert M. Weinstein, Mukesh Gupta, Kara S. Poole, Leon M. Bellan, Ethan S. Lippmann

**Affiliations:** Department of Chemical and Biomolecular Engineering, Vanderbilt University, TN, USA; Department of Biomedical Engineering, Vanderbilt University, Nashville, TN, USA; Department of Molecular Physiology and Biophysics, Vanderbilt University, Nashville, TN, USA; Interdisciplinary Materials Science Program, Vanderbilt University, Nashville, TN, USA; Department of Mechanical Engineering, Vanderbilt University, Nashville, TN, USA; Vanderbilt Brain Institute, Vanderbilt University, Nashville, TN, USA

**Keywords:** induced pluripotent stem cell, hydrogel, biomaterial, neuron, neural network

## Abstract

*In vitro* models of the human central nervous system (CNS), particularly those derived from induced pluripotent stem cells (iPSCs), are becoming increasingly recognized as useful complements to animal models for studying neurological diseases and developing therapeutic strategies. However, current 3D CNS models suffer from deficits that limit their research utility. Notably, it remains difficult to drive iPSC-derived neurons to a mature and synaptically connected state. Moreover, the most common extracellular matrices (ECMs) used to fabricate 3D CNS models are either difficult to pattern into complex structures due to their mechanical properties or lack appropriate bioinstructive cues. Here, we describe the functionalization of gelatin methacrylate (GelMA) with an N-cadherin extracellular peptide epitope to create a biomaterial termed GelMA-Cad. After photopolymerization, GelMA-Cad forms soft hydrogels that can maintain patterned architectures. The N-cadherin functionality promotes survival and maturation of iPSC-derived glutamatergic neurons into synaptically connected networks as determined by viral tracing and electrophysiology. Immunostaining reveals a pronounced increase in presynaptic and postsynaptic marker expression in GelMA-Cad relative to Matrigel, as well as extensive co-localization of these markers, thus highlighting the biological activity of the N-cadherin peptide. Overall, given its ability to enhance iPSC-derived neuron maturity and connectivity, GelMA-Cad should be broadly useful for *in vitro* studies of neural circuitry in health and disease.

## Introduction

Neurodegenerative diseases are estimated to affect 30 million people worldwide^[1]^. Aging is the number one risk factor for neurodegeneration, and disease incidence is expected to increase concurrently with an increasingly aged population worldwide. These diseases (e.g. Alzheimer’s disease, Parkinson’s disease, Huntington’s disease, Amyotrophic Lateral Sclerosis and Multiple Sclerosis) all have different region-specific presentation and modes of cell-cell communication, making it difficult to understand the mechanisms underlying their onset and propagation and thereby hampering development of effective treatments. As an example, clinical trials for Alzheimer’s disease have had a high failure rate since 2010^[2]^ and currently available drugs can alleviate symptoms but do not reverse neural tissue damage. These failures occur despite many candidate therapeutics having efficacy in various mouse models. As such, there has been a growing interest in developing human *in vitro* models for studying neurodegeneration^[3–5]^ and reducing the attrition rate in therapeutic development.

Recent advancements in 3D neural tissue models, particularly those constructed from human pluripotent stem cell (hPSC)-derived progenies (including human embryonic stem cells and induced pluripotent stem cells (iPSCs)), have generated much excitement for their ability to mimic the structure and function of human brain regions. Such models typically consist of neurons and varying mixtures of supporting cells (e.g. glia and vascular components) embedded in a hydrogel formed from extracellular matrix (ECM) components. Currently, a wide variety of ECMs are available for constructing neural tissues. Historically, the majority of early neural tissue models have utilized Matrigel, an ECM composite derived from Engelbreth-Holm-Swarm mouse sarcoma tumors that consists of many proteins (type IV collagen, laminin) and growth factors^[6]^. For example, 3D Matrigel scaffolds has been used to support differentiation of mouse embryonic stem cells to neural cells^[7]^. Matrigel is also the sole ECM utilized for more recently developed hPSC-derived brain organoids^[8,9]^, where the ECM scaffold supports the self-organization of the neuroepithelium to induce neuroepithelial buds and facilitates growth by providing a physical structure for cells to attach to and grow^[10]^. Other natural and synthetic materials have been developed for extended culture of hPSC-derived neural progenitor cells (NPCs) and neurons, including silk, collagen, hyaluronic acid (HA), elastin-like peptides, and polyethylene glycol (PEG)^[5,11–15]^. As these materials all allow for diffusion of essential nutrients and morphogens throughout the tissue constructs, they can be used to maintain NPCs and neuronal cultures for extended studies of differentiation and maturation, including axon formation, growth, and pruning^[12,16,17]^. Additionally, these platforms have demonstrated utility for assessing disease phenotypes when the hPSCs are sourced from patients that harbor genetic risk factors for each disorder^[11,13]^.

Despite progress towards fabricating complex neural tissue structures from hPSCs, existing ECMs have many shortcomings for practical neural cell culture. One limitation of existing ECM platforms is the lack of appropriate bioinstructive cues to promote cell-cell or cell-ECM interactions that facilitate neuronal maturation. Of the aforementioned materials, only Matrigel (e.g. laminin) and HA have physiological relevance to brain ECM; HA in particular has been shown to support hPSC-derived NPC maturation into neurons that exhibit enhanced neurite projection with synaptic vesicles and electrophysiological activity^[13]^. However, Matrigel and HA are difficult to handle due to their viscosity, and both materials have a very low elastic modulus, meaning they collapse under their own weight and cannot be molded into more complex structures. These factors limit the fabrication of topographic features, such as vasculature or perfusion channels. Synthetic hydrogels may overcome these issues by incorporating custom functional groups that enable tuning of mechanical and rheological properties, but they can be prohibitively difficult to fabricate and require extensive chemical modification to recapitulate tissue-specific biochemical cell-ECM interactions. Moreover, the majority of natural and synthetic ECMs are relatively expensive, which can further limit their widespread use.

Herein, we focused on engineering an ECM material to overcome some of these challenges. Our goal was to produce a material that would: (1) facilitate neuron survival and maturation within 3D tissue constructs through biophysical cues, (2) exhibit ideal mechanical properties to promote neuron outgrowth while also supporting micropatterned features, and (3) be relatively easy to synthesize, low cost, and therefore widely accessible. We ultimately produced a biomaterial termed GelMA-Cad, which consists of methacrylated gelatin (GelMA, a well-characterized biomaterial capable of being photopatterned^[18–20]^) conjugated with a peptide from an extracellular epitope of N-cadherin. Using this bioinstructive material, we form hydrogels with physiological stiffness that can not only maintain photopatterned features, but additionally facilitate iPSC-derived glutamatergic neuron survival and extension of neurite processes (as compared to GelMA, Matrigel, and various other negative controls). Moreover, relative to Matrigel, GelMA-Cad supports enhanced formation of synaptically connected neural networks, as measured by immunocytochemistry, electrophysiology, and viral synaptic tracing. Overall, we suggest that GelMA-Cad will aid the construction of 3D neural tissue models to study human disease biology and augment drug screening assays.

## Materials and methods

### Cell culture

CC3 iPSCs^[21]^ were maintained in E8 medium on standard tissue culture plastic plates coated with growth-factor reduced Matrigel (VWR). At 60-70% confluency, the cells were passaged using Versene (Thermo Fisher) as previously described^[22]^. Cortical glutamateric neurons were generated using a previously described protocol^[23]^ with some modifications. iPSCs were washed once with PBS and dissociated from the plates by incubation with Accutase (Thermo Fisher) for 3 minutes. After collection by centrifugation, cells were re-plated onto Matrigel-coated plates at a density of 2.5×10^5^ cells/cm^2^ in E8 medium containing 10 μM Y27632 (Tocris). The following day, the medium was switched to E6 medium^[22]^supplemented with 10 μM SB431542 (Tocris) and 0.4 μM LDN1931189 (Tocris) for 5 days to induce neuralization as previously described^[24]^. Over the next 5 days, the media was gradually transitioned from E6 medium to N2 Medium (DMEM/F12 basal medium (Thermo Fisher) containing 1X N2 supplement (Gibco), 10 μM SB431542, and 0.4 μM LDN193189)^[24]^. On the 11th day of the differentiation, the resultant neural progenitors were dissociated by incubation with Accutase for 1 hour and passaged onto Matrigel in Neural Maintenance Medium with 10 μM Y27632 at a cell density of 1×10^5^ cells/cm^2^. Neural Maintenance Medium was made by mixing a 1:1 ratio of N2 Medium and B27 Medium (Neurobasal Medium (Thermo Fisher) containing 200 mM Glutamax (Gibco) and 1X B27 (Gibco)). Cells received fresh Neural Maintenance Media every day for the next 20 days and a media change every 3-4 days afterwards. Neurons were used for experiments between days 70-100 of differentiation.

For the synaptic tracing experiments described below, a small population of neurons was also transduced with an adeno-associated virus (AAV) encoding EGFP under the control of the human synapsin promoter, which was a gift from Dr. Bryan Roth (Addgene plasmid #50465). Two weeks before the neurons were used, the cells were dissociated with Accutase and re-plated onto Matrigel-coated plates at a density of 2.5×10^5^ cells/cm^2^ in Neural Maintenance Media containing 10 μM Y27632. The following day, the media was replaced, and the AAV was added at a MOI of 5,000. Fresh media was added to the cells after 24 hours in order to remove any residual virus and normal media changes were resumed thereafter.

### GelMA synthesis and characterization

Methacrylated gelatin (GelMA) was synthesized as described previously^[25]^. Type A porcine skin gelatin (Sigma) was mixed at 10% (w/v) into DI water (sourced from an in-house Continental Modulab ModuPure reagent grade water system) at 60°C and stirred until fully dissolved. Methacrylic acid (MA) (Sigma) was slowly added to the gelatin solution and stirred at 50°C for 3 hours. The solution was then centrifuged at 3,500xg for 3 minutes and the supernatant was collected. Following a 5X dilution with additional warm (40°C) UltraPure water (Thermo Fisher) to stop the reaction, the mixture was dialyzed against DI water for 1 week at 37°C using 12–14 kDa cutoff dialysis tubing (Fisher) to remove salts and MA. The pH of the solution was then adjusted to 7.35-7.45 by adding HCl or NaOH as measured with a Thermo Fisher Scientific Orion Star pH meter. The resulting GelMA solution was lyophilized for 3 days using a Labconco lyophilizer and stored at −20°C.

### Peptide conjugation and characterization

Peptides were conjugated to GelMA as previously reported^[26]^ with slight modifications. Briefly, GelMA was reconstituted in triethanolamine (TEOA) buffer (Sigma) to create a 10% w/v solution and stirred at 37°C for 2 hours until fully dissolved. The pH of the solution was then adjusted to 8.0-8.5 using HCl or NaOH. Scrambled (Ac-AGVGDHIGC, to make GelMA-Scram) or N-Cadherin mimic (Ac-HAVDIGGGC, to make GelMA-Cad) peptides (GenScript) were added to the GelMA/TEOA buffer to form a 1% w/v solution. The cystine residue at the C-terminal end of the peptides permitted a Michael-type addition reaction with GelMA^[26]^. The solution was stirred at 37°C for 24 hours and then dialyzed against DI water using 6-8 kDa cutoff dialysis tubing (Spectrum) for 1 week at 37°C. The pH of the solution was then adjusted to 7.35-7.45 using HCl or NaOH, and the solution was lyophilized and stored at -20°C. Conjugation was routinely verified through ^1^H-NMR using a Bruker 500 Hz NMR spectrometer set to 37°C for the presence of the amino acid valine.

### Fourier-transform infrared spectroscopy

198 mg of potassium bromide (Sigma) was added to 2 mg of lyophilized gelatin, GelMA, GelMA-Cad, or GelMA-Scram and crushed using a mortar and pestle. The crushed samples were transferred to a 13 mm Specac evacuable pellet press die and compressed into a thin disc using a Specac manual hydraulic press. An additional disc was made using only potassium bromide for calibration. Samples were stored in a dry container overnight and analyzed the following day using a Bruker Tensor 27.

### Atomic force microscopy

GelMA, GelMA-Scram, and GelMA-Cad were reconstituted and polymerized into hydrogel discs as described in the cell seeding section below. A Bruker Dimension Icon Atomic Force Microscope was used to measure hydrogel stiffness. 0.01 N/m Novascan probes containing a 4.5 µm polystyrene bead (PT.PS.SN.4.5.CAL) were used to measure three distinct 5×5 µm areas of each hydrogel. Three hydrogel disc replicates of each sample were included for a total of 576 stiffness measurements per sample. For each individual force curve, a first order baseline correction was performed, and the Hertzian model was used to calculate Young’s modulus. For tool calibration, polyacrylamide hydrogels were prepared as previously reported^[27]^ and measured prior to GelMA and its derivatives.

### Scanning electron microscopy

Lyophilized GelMA, GelMA-Cad, and GelMA-Scram were reconstituted in PBS to form 10% (w/v) solutions with 0.05% LAP initiator (Sigma). 30 µL of each hydrogel solution was added to a Ted Pella pin mount and crosslinked by an 8 second exposure to a 25 mW/cm^2^ UV light using a ThorLabs UV Curing LED System. These pin mounts were stored in a Ted Pella mount storage tube and then placed in a -80°C freezer overnight. The following day, the samples were transferred to a Labconco lyophilizer for an additional 2 days and then stored at room temperature until used. To characterize the internal microscructures of GelMA, GelMA-Cad and GelMA-Scram, the dried samples were observed using a scanning electron microscope (Zeiss Merlin) at an accelerating voltage of 2 kV. ImageJ software was used to quantify pore sizes, where the mean diameter of each pore was considered the average pore size.

### Fabrication and seeding of hydrogel scaffolds

GelMA, GelMA-Scram and GelMA-Cad were reconstituted in Neuron Maintenance Media to make a 10% (w/v) solution with 0.05% LAP initiator. iPSC-derived neurons were detached from 12-well plates via a 5 minute incubation with Accutase and centrifuged for collection. Unless otherwise stated, neurons were mixed with reconstituted hydrogel/initiator solution to achieve a density of 2×10^5^ cells/mL. For some experiments, GelMA was mixed with soluble peptide rather than via covalent coupling; here, soluble peptides were reconstituted in DMSO to create a 10 mg/mL solution, and then the peptides were added to the GelMA/initiator/neuron solution to achieve a 50 µg/mL peptide concentration. Once the solutions were prepared, they were mixed thoroughly with a P1000 pipette to break up any cell clumps. Next, 100 µL of the cell suspension was added to RainX-treated glass slides and covered with 12 mm diameter coverslips (Carolina) to form discs. These discs were then exposed to 25 mW/cm^2^ UV light for 8 seconds and set aside for 10 minutes at room temperature. Hydrogel discs were then removed from the glass slides and transferred to a 12-well plate with 1 mL of Neural Maintenance Media per well.

To embed neurons in Matrigel, 1 mL Matrigel aliquots were thawed on ice. Once thawed, the neurons were embedded at the same cell density as described above, and 100 µl of the solution was added directly onto the coverslips in a 12-well plate. The plate containing the Matrigel discs was placed in an incubator at 37°C to crosslink for 30 minutes. After the Matrigel was fully crosslinked, 1 mL of Neural Maintenance Media was added to each of the wells. For all conditions, media was replaced twice a week until cells were used.

### Live/dead cell imaging

To assess long term cell viability, hydrogel discs were incubated with CytoCalcein™ Violet 450 (AAT Bioquest) and propidium iodide (PI, Thermo Fisher) for one hour. The hydrogel discs were imaged using a Zeiss 710 confocal microscope and cell viability was quantified using ImageJ. Following imaging, 1 mL of Neural Maintenance Media was added to each well in order to dilute any remaining Calcein/PI from the hydrogels.

### Neurite projection quantification

Raw data were exported in 16-bit TIF format and imported into Matlab 2017 for quantification using a custom image analysis script. Briefly, images were smoothed using a 3×3 pixel smoothing filter to mitigate image noise, and in-focus neurite segments were identified by isolating regions at least 5% brighter than the mean pixel intensity in the surrounding 50-pixel radius^[28]^. Cell bodies and neurites were distinguished by successive erosion of the resulting binary mass. The erosion radius at which the total cell mass declined most steeply was used to define the radius required to erode neurites while sparing cell bodies. Following segmentation of neurites and cell bodies, algorithms previously developed for analysis of mitochondrial networks^[29]^ were used to measure the average length and width of each neurite segment.

### Synaptic tracing

Hydrogel discs were fabricated as described above. Prior to crosslinking (of GelMA-Cad) or gelation (of Matrigel), neurons transduced with synapsin-driven EGFP were dissociated from plates via a 5 minute incubation with Accutase and then added to the center of the hydrogel disc at a density of 2×10^3^ cells/mL (as shown in **Figure 6**). After crosslinking or gelation, the hydrogel discs were placed in 1 mL of Neural Maintenance Media and stored in an incubator at 37°C until imaged. For all conditions, the media was replaced twice a week. The formation of synaptic connections was visualized by the spread of EGFP fluorescence across each hydrogel using a Zeiss LSM 710 confocal microscope.

### Immunofluorescence

After 2 weeks of culture, neurons embedded in hydrogels were fixed in 4% PFA (Sigma) for 20 minutes and then washed 3 times with PBS. A solution of 5% goat serum and 0.03% Triton X-100 (Thermo Fisher) was then added to the hydrogels overnight on a rocking platform at room temperature. The hydrogels were then incubated overnight with DAPI and a combination of the following fluorescently conjugated primary antibodies: βIII tubulin Alexa Fluor 647 (Abcam ab190575), PSD-95 Alexa Fluor 488 (Novus Biologicals NB300556AF488), and/or synaptophysin Alexa Fluor 555 (Abcam ab206870). Hydrogels were then imaged using a 40x objective on a Zeiss LSM 710 confocal microscope. The number of PSD-95 and synaptophysin puncta was quantified using the cell counter plugin on ImageJ. Colocalization of these two markers was quantified using Zeiss Zen Black software.

### Electrophysiology

Neurons embedded in GelMA-Cad or Matrigel hydrogels were recorded in a bath consisting of 140 mM NaCl, 2.8 mM KCl, 2 mM CaCl_2_, 2 mM MgCl_2_, 10 mM HEPES, and 10 mM D-glucose. Sharp glass microelectrodes were prepared from borosilicate glass with a Sutter P97 pipette puller and filled with extracellular solution to reach a resistance of 6−8 MΩ. The recording electrode was placed near the edge of the hydrogel disc. Whole-cell patch clamp recordings were performed in a recording chamber placed on the stage of a Zeiss Axioscope upright microscope. Current clamp experiments were performed with an Axon Multiclamp 700A amplifier. Data recording and analysis were performed with Axon pClamp software.

## Results

### Synthesis and characterization of GelMA functionalized with N-cadherin peptide

GelMA was chosen as a base material due to its ease of handling and robust mechanical properties (after crosslinking) compared to ECMs such as Matrigel and HA. Meanwhile, N-cadherin functionality was chosen for the role of this cell adhesion molecule in neurite growth during neurogenesis^[30–35]^. The extracellular peptide epitope of N-cadherin chosen for this study has previously been used to functionalize methacrylated HA in order to support chondrogenesis from mesenchymal stem cells^[26]^, but 3D scaffolds fabricated with this peptide have not been used to support neural cultures. To generate the GelMA-Cad scaffold, porcine gelatin was first functionalized with methacrylic anhydride in order to create the GelMA backbone that could be crosslinked when exposed to the photoinitiator LAP and UV light (**Figure 1**)^[36]^. This modification was confirmed through the presence of methacrylic side chain protons (∼5.45 and 5.7 ppm) using ^1^H-NMR (**Figure 2B**). GelMA was then functionalized with the extracellular epitope of N-cadherin (HAVDIGGGC) or an N-cadherin-scrambled peptide (AGVGDHIGC), which is referred to as GelMA-Scram. The conjugation of these peptides to the scaffold was also confirmed via ^1^H NMR through the presence of valine protons (∼3.5 ppm), which are not present in the gelatin or GelMA spectra (**Figure 2B**). Additionally, Fourier-transform infrared spectroscopy (FTIR) was employed to further validate successful functionalization. The FTIR transmittance spectra showed a noticeable decrease in PO_4_ peaks (1000cm^-1^) and amide peaks I, II, III (1640, 1540, and 1250 cm^-1^, respectively) in GelMA-Cad and GelMA-Scram samples compared to GelMA (**Figure 2C**), likely due to peptide conjugation. Collectively, these data suggest GelMA was properly synthesized and functionalized.

**Figure 1:**
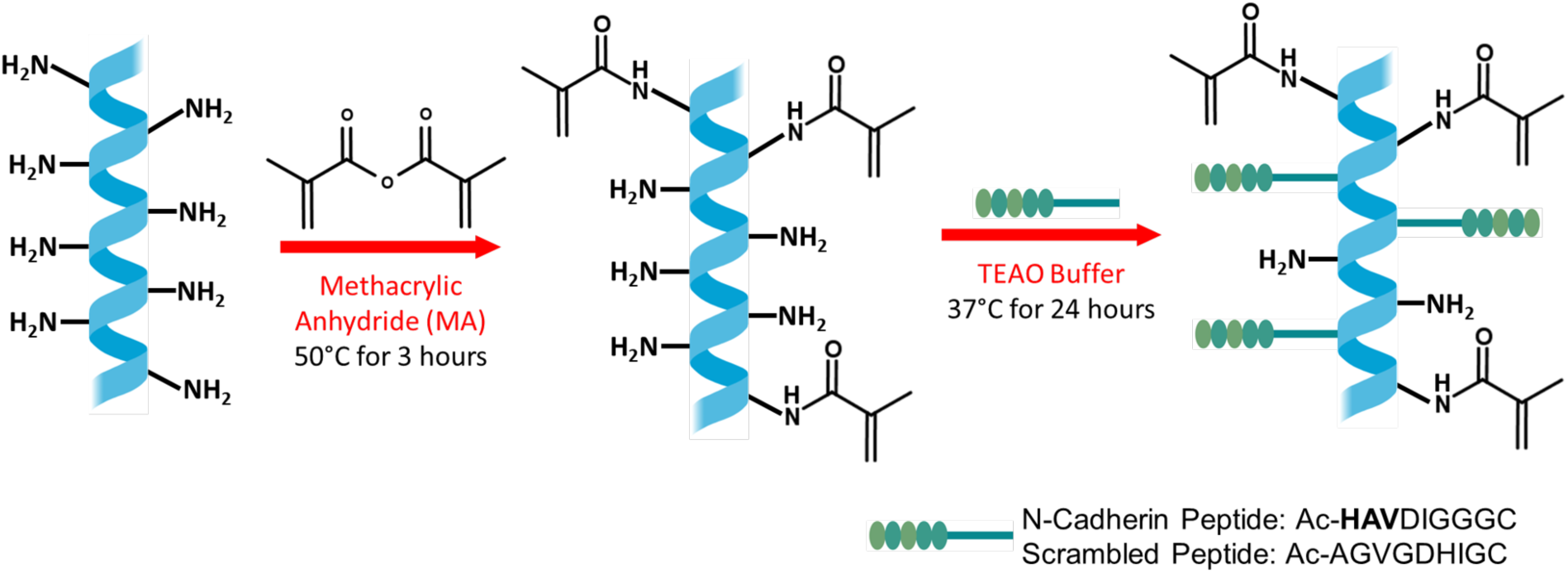
Schematic illustration of GelMA synthesis and N-cadherin peptide conjugation. The conventional method for synthesizing GelMA uses methacrylic anhydride to introduce a methacryloyl substitution group on the reactive primary amine group of amino acid residues. GelMA was then dissolved in TEAO buffer with the N-cadherin peptide for Michael-type addition to the reactive primary amine group of the amino acid.

**Figure 2:**
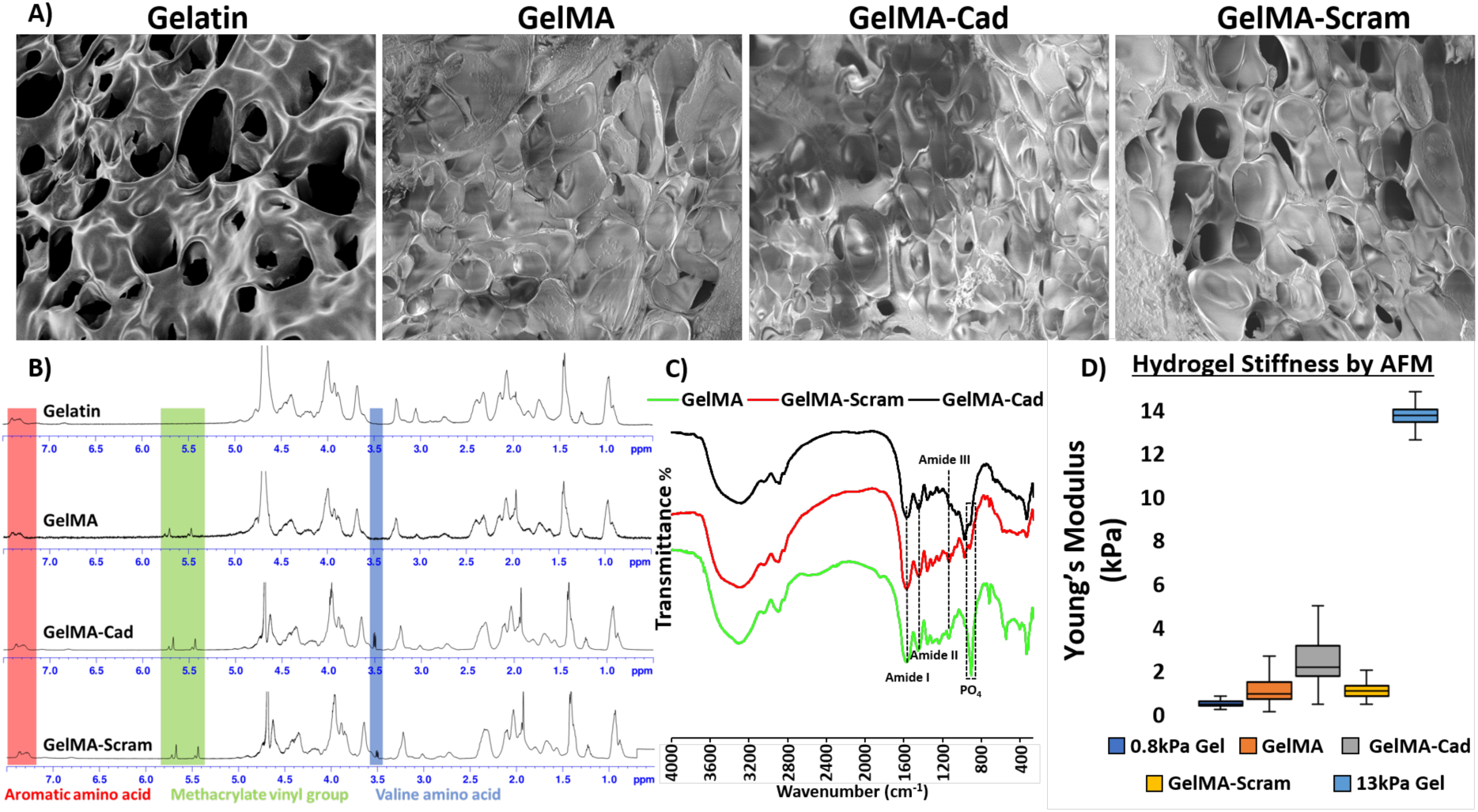
Assessment of biomaterial functionalization and physical properties of polymerized hydrogels. **(A)** SEM images of hydrogels fabricated from gelatin, GelMA, GelMA-Cad, and GelMA-Scram. **(B)** NMR spectra of gelatin, GelMA, GelMA-Cad, and GelMA-Scram. Successful conjugation of methacrylic anhydride to the backbone of gelatin was assessed by peaks at 5.5 and 5.7 ppm, and N-cadherin/Scram peptide addition was assessed by the valine peak at 3.5 ppm. **(C)** FTIR spectra was used to confirm conjugation of the peptide to the backbone of GelMA due to decrease in the following relevant bands: 1000 cm^−1^ (PO_4_ stretching) and 1250 cm^−1^, 1540 cm^−1^, and 1640 cm^−1^ (NH bending). **(D)** AFM measurements of Young’s modulus values for GelMA, GelMA-Cad, GelMA-Scram. Data are presented as mean ± S.D. from 3 independently fabricated hydrogels, where three locations were sampled on each hydrogel as described in the methods.

### Mechanical and physical properties of crosslinked hydrogels

In order to determine the stiffness of GelMA, GelMA-Cad, and GelMA-Scram, atomic force microscopy (AFM) was performed. 0.8 kPa and 13 kPa polyacrylamide hydrogels were produced and measured by AFM^[27]^ to validate that the tool was properly calibrated (**Figure 2D**). After crosslinking with LAP and UV light, GelMA, GelMA-Cad, and GelMA-Scram exhibited stiffness values of approximately 1-5 kPa (**Figure 2D**), which resembles the stiffness of native brain tissue^[37]^. Despite its relatively low elastic modulus, GelMA-Cad is stiff enough to maintain patterned architectures: when it was crosslinked around silicone tubing, followed by manual extraction of the tubing, a straight, a perfusable channel remained in the GelMA-Cad (**Supplemental Figure 1A**), whereas Matrigel collapses and the perfusion channel does not remain patent (**Supplemental Figure 1B**). Thus, similar to GelMA^[36,38]^, GelMA-Cad can be patterned into more complex structures.

The microstructure of the hydrogels was characterized by scanning electron microscopy (SEM). Porous network structures are commonly observed in hydrogels and are important for nutrient diffusion, cell integration and removal of waste products, and the degree of chemical substitution has an inverse relation to pore size upon crosslinking ^[39,40]^. The average pore size diameter of GelMA, GelMA-Cad, and GelMA-Scram were measured at 42.8 ± 0.2, 43.1 ± 0.2, and 42.4 ± 0.2 µm, respectively (**Figure 2A**). These measurements confirm that the hydrogels all have similar physical and mechanical properties, such that differences in neuron behavior can likely be attributed to bioinstructive cues.

### GelMA-Cad hydrogels support survival and outgrowth of iPSC-derived neurons

To assess the ability of hydrogels to support human neuron survival and outgrowth, human iPSCs were differentiated into cortical glutamatergic neurons and cultured for 70-100 days before use. These neurons were then dissociated into single-cell suspensions and embedded into Matrigel, GelMA-Cad, GelMA-Scram, or GelMA. As a negative control for physical conjugation of peptides to the hydrogels, neurons were also embedded in GelMA with either soluble N-cadherin peptide or soluble scrambled peptide. Using calcein and propidium iodide dyes to mark live and dead cells, respectively, we determined that neurons embedded in GelMA and GelMA-Scram (both conjugated and soluble peptide), as well as Matrigel with the soluble peptide, died within 4 days (**Figure 3A, Figure 3B, and Supplemental Figure 2**). Meanwhile, neurons in conjugated GelMA-Cad and Matrigel exhibited viability of 90.2 ± 1.3% and 86.3 ± 2.2% after 2 days, respectively. After 3 days, neurons in conjugated GelMA-Cad exhibited viability of 96.7 ± 1.2% while viability in Matrigel decreased slightly to 80 ± 1.3%. After 5 days, viability remained relatively constant (96.7 ± 1.1% in conjugated GelMA-Cad versus 82.3 ± 1.9% in Matrigel). By day 10, viability in conjugated GelMA-Cad continued to remain constant at 96.7 ± 1.6% whereas viability in Matrigel again decreased slightly to 76.7 ± 0.8%. Next, we monitored neurite projections from neurons embedded in either Matrigel or conjugated GelMA-Cad (referred to solely as GelMA-Cad from hereon) after 5 and 10 days using calcein. Neurite length and width are frequently employed as measures of neuron health and connectivity^[41–44]^, and so we quantified Z-stack images of neurites using a custom Matlab script (**Figure 4A-H**). On day 5, relative to Matrigel, neurons embedded in GelMA-Cad exhibited significantly higher average neurite length (28.9 ± 1.6 µm vs 14.1 ± 2.6 µm; p<0.05), whereas average neurite width was not significantly different between GelMA-Cad and Matrigel (4.0 ± 0.2 µm vs 3.7 ± 0.2 µm) (**Figure 4I-J**).

**Figure 3:**
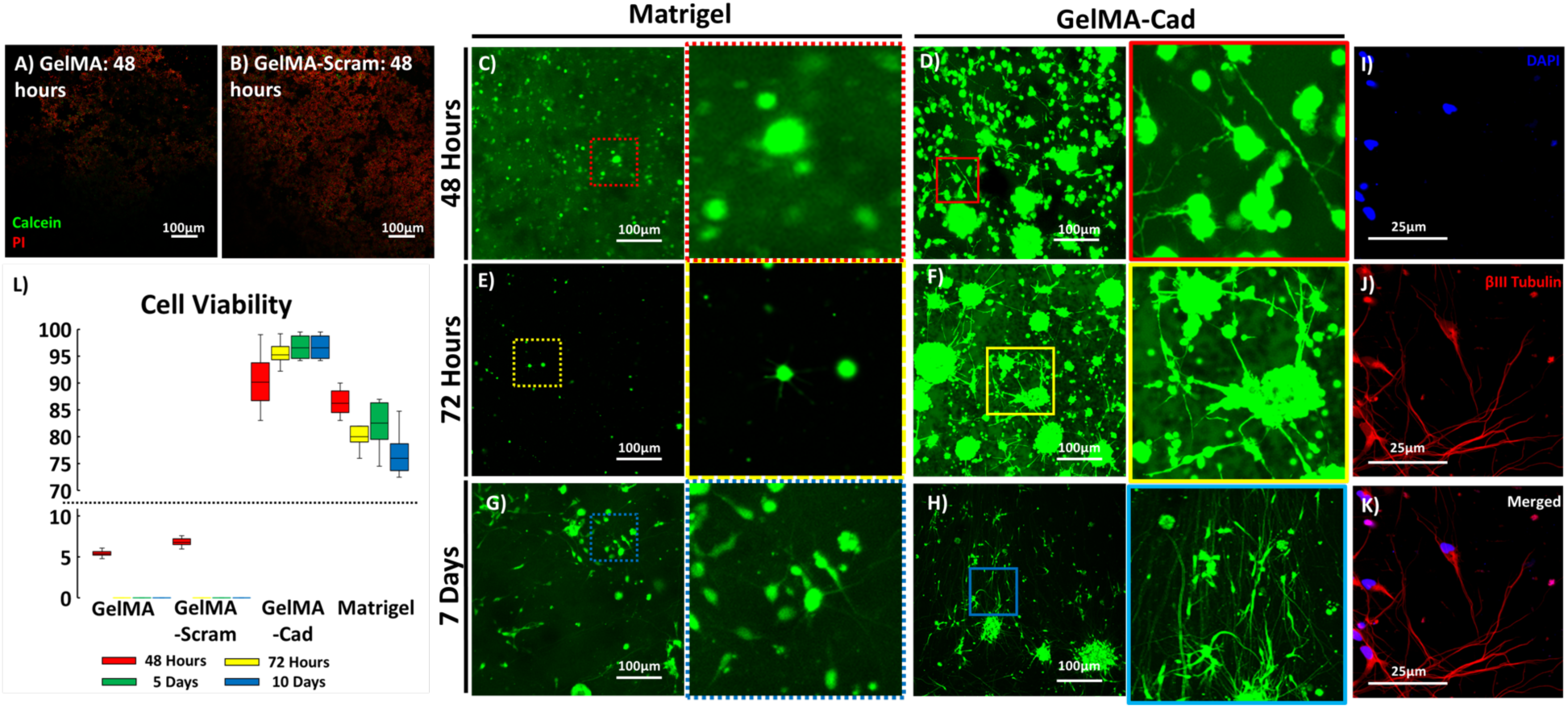
Live/dead staining of iPSC-derived neurons embedded in various hydrogels. For panels A-H, cells were labeled with calcein (green) to visualize live cells and propidium iodide (PI, red) to visualize dead cells. For panels A-B, both calcein and PI staining are shown to highlight dead cells. For panels C-H, only calcein is shown to highlight neuron morphology in GelMA-Cad, and insets are provided for higher magnification. Full quantification of viability is shown in panel L. **(A)** Neurons in GelMA 48 hours after embedding. **(B)** Neurons in GelMA-Scram 48 hours after embedding. **(C-D)** Neurons in Matrigel or GelMA-Cad 48 hours after embedding. **(E-F)** Neurons in Matrigel or GelMA-Cad 72 hours after embedding. **(G-H)** Neuron in Matrigel or GelMA-Cad 7 days after embedding. **(I-K)** Neurons in GelMA-Cad were immunolabeled 7 days after embedding for βIII tubulin (red) to confirm identity. **(L)** Cell viability is presented for various time points as mean ± S.D. from 3 biological replicates, with 5 images assessed per replicate.

**Figure 4:**
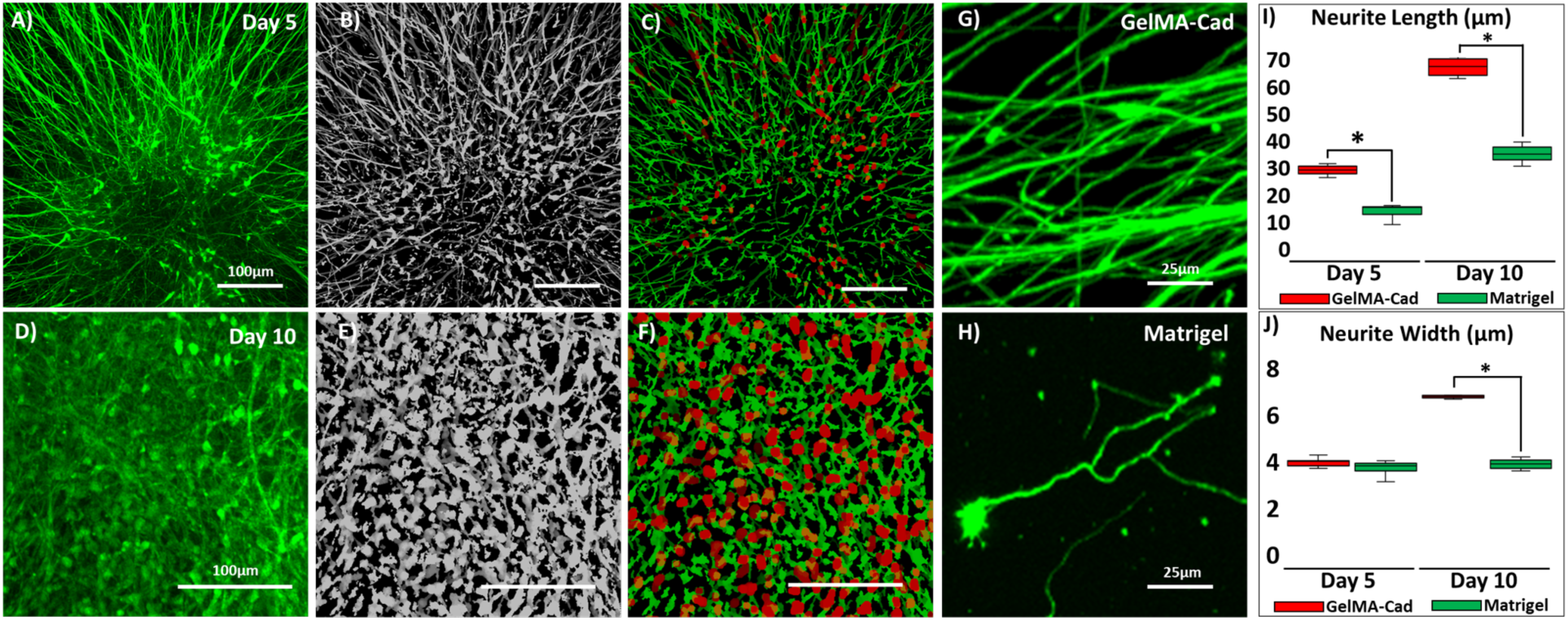
Quantification of neurites in iPSC-derived neurons embedded in Matrigel and GelMA-Cad. Panels A-C demonstrate the quantification of neurites in GelMA-Cad on day 5, and panels D-E demonstrate the quantification of neurites in GelMA-Cad on day 10. Neurons are stained with calcein (green) and imaged with a confocal microscope **(A, D)**. Using custom Matlab code, a mask is applied **(B, E)** and cell soma and neurites are identified **(C,F)**, where red corresponds to the soma and green corresponds to neurite extensions, which can then be measured and averaged across an image. **(G-H)** Example of high-resolution images of neurites in GelMA-Cad and Matrigel, where differences in neurite length and thickness can be observed. **(I-J)** Full quantification of neurite length and width. Data are presented as mean ± S.D. from 7 biological replicates, with 4 images quantified per replicate. Statistical significance was calculated using the student’s unpaired t-test (*, p<0.05).

However, on day 10, relative to Matrigel, neurons in GelMA-Cad exhibited significantly higher average neurite length (67.2 ± 3.2 µm vs 35.3 ± 7.1 µm; p<0.05) and average neurite width (6.8 ± 0.2 µm vs 3.9 ± 0.2 µm; p<0.05) (**Figure 4I-J**). These results demonstrate GelMA-Cad is an effective hydrogel for enhancing survival and maturation of human iPSC-derived neurons by morphometric parameters.

### iPSC-derived neurons form synaptically connected networks in GelMA-Cad hydrogels

The increased length and diameter of neurons in GelMA-Cad suggests improved functional properties, which we sought to validate with additional metrics including immunostaining, electrophysiological recordings, and viral synaptic tracing. First, we fixed and immunostained embedded neurons for synaptophysin (a presynaptic terminal marker) and PSD-95 (a postsynaptic terminal marker). Neurons embedded in GelMA-Cad expressed both markers 21 days after embedding (average of 492 synaptophysin puncta and 423 PSD-95 puncta per 75 µm^3^), and there was an average of 87.3 ± 1.3% co-localization, which indicates the formation of an active synapse (**Figure 5A**). Neurons embedded in Matrigel had substantially lower expression of synaptophysin and PSD-95 (average of 82 puncta and 28 puncta per 75 µm^3^, respectively), with only 13.3 ± 3.3% colocalization of the presynaptic and postsynaptic markers (**Figure 5A**), indicating a substantially lower number of prospective synapses. Next, to assess synaptic connectivity, we measured electrical activity of the embedded neurons through patch clamping. Action potentials were readily measured within neurons embedded in GelMA-Cad (**Figure 5B, red line**), but only minimal activity was observed in Matrigel-embedded neurons (**Figure 5B, black line**), thus providing evidence that the N-cadherin peptide improves functional maturity.

**Figure 5:**
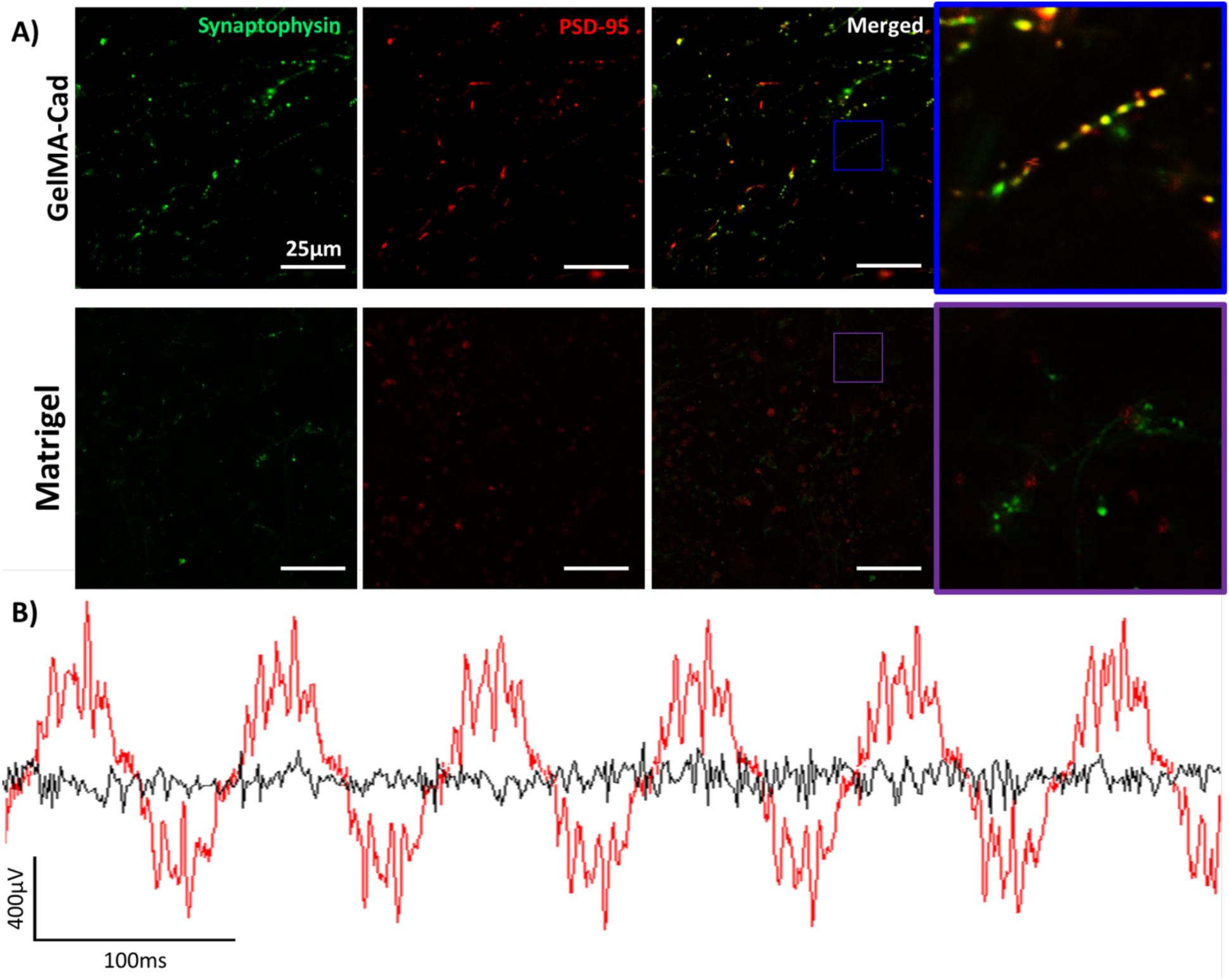
Assessment of synaptic connectivity of iPSC-derived neurons in Matrigel or GelMA-Cad by immunostaining and electrophysiology. **(A)** Immunostaining of synaptophysin (green) and PSD-95 (red) in neurons that were embedded in each hydrogel for 21 days. An inset is provided to highlight pronounced differences. 10 images from 3 biological replicates were used for absolute quantification of expression and percent co-localization. **(B)** Electrical activity in neurons embedded in GelMA-Cad (red) and Matrigel (black) for 21 days. Voltage traces are representative of 5 biological measurements.

**Figure 6:**
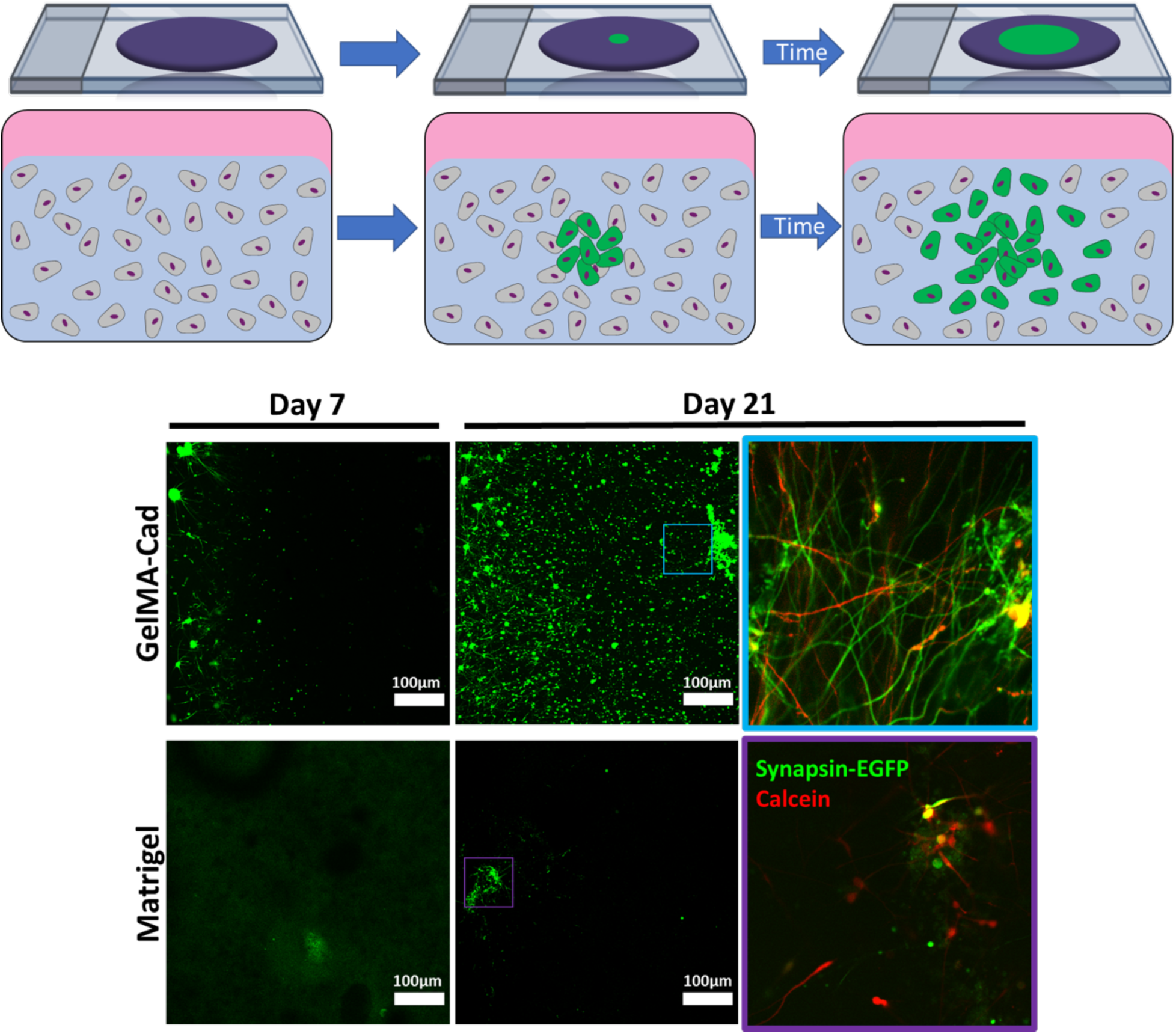
Assessment of synaptic connectivity of iPSC-derived neurons in Matrigel or GelMA-Cad by viral tracing. The schematic depicts the experimental approach, where wild-type neurons were uniformly mixed in a hydrogel and a small population of AAV-transduced neurons were injected into the center. EGFP was then imaged at day 7 and day 21. Calcein (red) was added at day 21 to verify cell viability as highlighted by the insets. The images from these experiments are representative of 6 biological replicates that confirmed the transmission of EGFP in GelMA-Cad but not Matrigel.

Last, to assess widespread neural network formation, we conducted synaptic tracing experiments by transducing iPSC-derived neurons with an adeno-associated virus (AAV) encoding EGFP under the control of human synapsin promoter (where synapsin is a presynaptic terminal marker). Wild-type neurons were mixed with hydrogel precursor, and prior to crosslinking the hydrogels, a small population of AAV-transduced neurons (1:100 ratio of transduced to non-transduced neurons) were injected into the center (**Figure 6**). The spread of the EGFP signal could thus be monitored over time to elucidate the degree of neural network formation across the hydrogel. Limited EGFP spread was observed after 7 days in both hydrogels, which is consistent with **Figure 4** demonstrating that neurite length and width are still increasing at this early time point. However, after 21 days, EGFP had propagated to virtually every neuron within the GelMA-Cad hydrogels, whereas sparse EGFP spread was observed in Matrigel (**Figure 6**). Calcein dye was added to each hydrogel to show that the neurons in Matrigel were alive but not synaptically connected. Therefore, only neurons in the GelMA-Cad hydrogels were able to propagate the virus through functional synapses across the entire tissue construct. Overall, these data strongly suggest that GelMA-Cad facilitates the maturation of iPSC-derived neurons on a functional level.

## Discussion

In native tissues, cells are surrounded by a complex ECM that presents both physical and biochemical cues for tissue development, migration and proliferation. In the brain, neurons are encased in a soft ECM comprised of various proteins and glycosaminoglycans (GAGs). Many natural and synthetic ECMs, as described earlier, have been used to support 3D *in vitro* cultures of neural cells. Several of these studies have focused on the effect of ECM stiffness and composition on the differentiation of NPCs to terminally differentiated progenies, but relatively fewer have examined functional maturation and integration of neurons, particularly those derived from hPSCs. hPSC-derived neurons are notoriously difficult to mature in 2D without extended culture times or co-culture with astrocytes, though more success has been observed in 3D assembly. In particular, hydrogels comprised of methacrylated HA can facilitate the maturation of iPSC-derived neurons to an electrophysiologically active state, which is perhaps unsurprising given that HA is the most abundant GAG in brain tissue. Yet, as mentioned previously, HA has limitations that make it less attractive for either large-scale work or applications requiring patterned structures.

Our approach to these challenges was to use a relatively simple ECM material and append it with a biophysical cue to mimic cell-cell communication in the developing and adult brain. N-cadherin was chosen for several reasons. From a biomaterials perspective, the N-cadherin extracellular epitope had already been incorporated into HA gels and shown to be bioactive for cartilage tissue engineering, so it was straightforward to incorporate this epitope into GelMA. Moreover, tethering of an N-cadherin peptide to a solid support had previously been shown to improve neuronal survival and neurite outgrowth of stem cell-derived neurons in 2D cultures^[30,45–47]^, suggesting relevance for our work in 3D scaffolds. From a biochemical perspective, N-cadherin expression is important for outgrowth of neurites on astrocyte monolayers^[48]^, and given that hPSC-derived neurons are typically cultured on 2D monolayers of astrocytes to facilitate electrophysiological maturation, we suspected that GelMA-Cad could partially replace the synaptogenic role of astrocyte co-culture. Cadherins also regulate dendritic spine morphology^[49]^ and N-cadherin-mediated interactions are specifically required for maintenance of dendrites^[50]^, which is particularly interesting because we observed an especially pronounced increase in the expression of postsynaptic terminal markers on neurons in GelMA-Cad relative to Matrigel.

It is also possible that the gelatin backbone of GelMA-Cad augments neuron health and survival. Gelatin-based hydrogels can be neuroprotective and promote neurite outgrowth through integrin activation and integrin-dependent MAPK signaling^[50]^. Gelatin also improves *in vivo* outcomes in dissected nerve regeneration, particularly through increased axonal migration^[51–53]^. Recent work has further shown that GelMA alone can support dissociated primary mouse neurons to create a synaptically connected neural network on the micron scale^[54]^. This finding contrasts somewhat with our observations of neuron death in GelMA and GelMA-Scram. However, beyond the obvious species differences, we note that this prior study utilized neurons that had already matured *in vivo*. Thus, the primary neurons may have been competent enough to form neural networks in GelMA because they already expressed the requisite synaptic machinery, rather than requiring *de novo* expression of this machinery in hPSC-derived neurons developing from a more embryonic-like state.

Overall, our work provides an exciting scaffolding material for future efforts devoted to assembling complex neurovascular tissue structures. If N-cadherin signaling is a general mechanism for synaptic integration, we speculate that multiple iPSC-derived neuronal cell types could be patterned into complex architectures that would be subsequently directed by the ECM to form relevant neural circuitry. The inclusion of iPSC-derived astrocytes and microglia could further be used to study synaptic pruning and remodeling under healthy and diseased conditions. Then, given the mechanical integrity of GelMA-Cad hydrogels, various micropatterning approaches could be used to generate perfusable channels seeded with endothelial and mural cells, ultimately resulting in functional vasculature throughout the human neural networks. The bioactivity of GelMA-Cad, coupled to its relative low cost and ease of synthesis, should make it attractive for the aforementioned applications, as well as other situations throughout bioengineering and neuroscience.

## Acknowledgments

The authors would like to thank Dr. Anthony Hmelo and the Vanderbilt Institute of Nanoscale Science and Engineering (VINSE) for assistance with atomic force microscopy and scanning electron microscopy. The authors would also like to thank Dr. Laura Dugan and Jacob Ruden for use of electrophysiology equipment. Funding for this work was provided by a Ben Barres Early Career Acceleration Award from the Chan Zuckerberg Initiative (grant 2018-191850 to ESL), NSF grants 1846860 and 1706155 (to ESL) and 1506717 and 1462866 (to LMB), a NARSAD Young Investigator Award from the Brain and Behavior Research Foundation (grant 25177 to ESL), the donors of Alzheimer’s Disease Research which is a program of the BrightFocus Foundation (grant A20170945 to ESL), NIH grant R00EB013630 (to LMB), and NIH grant 5P30 DK020593 (pilot award to ESL). BJO was partially supported by the NIH-funded Vanderbilt Interdisciplinary Training Program in Alzheimer’s Disease (T32 AG058524). KMB was partially supported by the NIH-funded Training Program in Environmental Toxicology (T32 ES007028). PMM was supported by an NIH NRSA fellowship (F32 DK120104).

## References

[1] S. Vanni, A. Colini Baldeschi, M. Zattoni, G. Legname, J. Neurosci. Res. 2019.

[2] D. Mehta, R. Jackson, G. Paul, J. Shi, M. Sabbagh, Expert Opin. Investig. Drugs 2017, 26, 735.

[3] J. A. Bolker, BioEssays 2017, 39.

[4] D. Zhang, M. Pekkanen-Mattila, M. Shahsavani, A. Falk, A. I. Teixeira, A. Herland, Biomaterials 2014, 35, 1420.

[5] P. Zhuang, A. X. Sun, J. An, C. K. Chua, S. Y. Chew, Biomaterials 2018, 154, 113.

[6] C. S. Hughes, L. M. Postovit, G. A. Lajoie, Proteomics 2010, 10, 1886.

[7] C. R. Kothapalli, R. D. Kamm, Biomaterials 2013, 34, 5995.

[8] A. Fatehullah, S. H. Tan, N. Barker, Nat. Cell Biol. 2016, 18, 246.

[9] M. A. Lancaster, M. Renner, C. A. Martin, D. Wenzel, L. S. Bicknell, M. E. Hurles, T. Homfray, J. M. Penninger, A. P. Jackson, J. A. Knoblich, Nature 2013, 501, 373.

[10] H. Wang, Front. Synaptic Neurosci. 2018, 10, 15.

[11] W. L. Cantley, C. Du, S. Lomoio, T. Depalma, E. Peirent, D. Kleinknecht, M. Hunter, M. D. Tang-Schomer, G. Tesco, D. L. Kaplan, ACS Biomater. Sci. Eng. 2018, 4, 4278.

[12] K. Pietrucha, M. Zychowicz, M. Podobinska, L. Buzanska, Folia Neuropathol. 2017, 55, 110.

[13] Z.-N. Zhang, B. C. Freitas, H. Qian, J. Lux, A. Acab, C. A. Trujillo, R. H. Herai, V. A. Nguyen Huu, J. H. Wen, S. Joshi-Barr, J. V. Karpiak, A. J. Engler, X.-D. Fu, A. R. Muotri, A. Almutairi, Proc. Natl. Acad. Sci. 2016, 113, 3185.

[14] C. Barry, M. T. Schmitz, N. E. Propson, Z. Hou, J. Zhang, B. K. Nguyen, J. M. Bolin, P. Jiang, B. E. McIntosh, M. D. Probasco, S. Swanson, R. Stewart, J. A. Thomson, M. P. Schwartz, W. L. Murphy, Exp. Biol. Med. 2017, 242, 1679.

[15] C. M. Madl, B. L. Lesavage, R. E. Dewi, C. B. Dinh, R. S. Stowers, M. Khariton, K. J. Lampe, D. Nguyen, O. Chaudhuri, A. Enejder, S. C. Heilshorn, Nat. Mater. 2017, 16, 1233.

[16] M. Georgiou, S. C. J. Bunting, H. A. Davies, A. J. Loughlin, J. P. Golding, J. B. Phillips, Biomaterials 2013, 34, 7335.

[17] K. Chwalek, D. Sood, W. L. Cantley, J. D. White, M. Tang-Schomer, D. L. Kaplan, J. Vis. Exp. 2015, 105, e52970.

[18] X. Zhou, H. Cui, M. Nowicki, S. Miao, S. J. Lee, F. Masood, B. T. Harris, L. G. Zhang, ACS Appl. Mater. Interfaces 2018, 10, 8993.

[19] S. K. Davey, A. Aung, G. Agrawal, H. L. Lim, M. Kar, S. Varghese, Tissue Eng. Part C Methods 2015, 21, 1188.

[20] J. W. Nichol, S. T. Koshy, H. Bae, C. M. Hwang, S. Yamanlar, A. Khademhosseini, Biomaterials 2010, 31, 5536.

[21] K. K. Kumar, E. W. Lowe, A. A. Aboud, M. D. Neely, R. Redha, J. A. Bauer, M. Odak, C. D. Weaver, J. Meiler, M. Aschner, A. B. Bowman, Sci. Rep. 2014, 28, 6801.

[22] E. S. Lippmann, M. C. Estevez-Silva, R. S. Ashton, Stem Cells 2014, 32, 1032.

[23] Y. Shi, P. Kirwan, F. J. Livesey, Nat. Protoc. 2012, 7, 1836.

[24] S. M. Chambers, C. A. Fasano, E. P. Papapetrou, M. Tomishima, M. Sadelain, L. Studer, Nat. Biotechnol. 2009, 27, 275.

[25] D. Loessner, C. Meinert, E. Kaemmerer, L. C. Martine, K. Yue, P. A. Levett, T. J. Klein, F. P. W. Melchels, A. Khademhosseini, D. W. Hutmacher, Nat. Protoc. 2016, 11, 727.

[26] L. Bian, M. Guvendiren, R. L. Mauck, J. A. Burdick, Proc. Natl. Acad. Sci. 2013, 110, 10117.

[27] K. M. Stroka, H. Aranda-Espinoza, Blood 2011, 118, 1632.

[28] P. M. McClatchey, N. A. Mignemi, Z. Xu, I. M. Williams, J. E. B. Reusch, O. P. McGuinness, D. H. Wasserman, Microcirc. 2018, 25, e12482.

[29] P. M. McClatchey, A. C. Keller, R. Bouchard, L. A. Knaub, J. E. B. Reusch, Mitochondrion 2016, 26, 58.

[30] A. Haque, N. Adnan, A. Motazedian, F. Akter, S. Hossain, K. Kutsuzawa, K. Nag, E. Kobatake, T. Akaike, PLOS ONE 2015, 10, e0135170.

[31] M. Kadowaki, S. Nakamura, O. Machon, S. Krauss, G. L. Radice, M. Takeichi, Dev. Biol. 2007, 304, 22.

[32] J. D. Jontes, Cold Spring Harb. Perspect. Biol. 2018, 10, a029306.

[33] C. Xu, Y. Funahashi, T. Watanabe, T. Takano, S. Nakamuta, T. Namba, K. Kaibuchi, J. Neurosci. 2015, 35, 14517.

[34] M. Shikanai, K. Nakajima, T. Kawauchi, Commun. Integr. Biol. 2011, 4, 326.

[35] Y. Matsunaga, M. Noda, H. Murakawa, K. Hayashi, A. Nagasaka, S. Inoue, T. Miyata, T. Miura, K. Kubo, K. Nakajima, Proc Natl Acad Sci 2017, 114, 2048.

[36] D. Loessner, C. Meinert, E. Kaemmerer, L. C. Martine, K. Yue, P. A. Levett, T. J. Klein, F. P. W. Melchels, A. Khademhosseini, D. W. Hutmacher, Nat. Protoc. 2016, 11, 727.

[37] S. Budday, R. Nay, R. de Rooij, P. Steinmann, T. Wyrobek, T. C. Ovaert, E. Kuhl, J. Mech. Behav. Biomed. Mater. 2015, 46, 318.

[38] L. E. Bertassoni, M. Cecconi, V. Manoharan, M. Nikkhah, J. Hjortnaes, A. L. Cristino, G. Barabaschi, D. Demarchi, M. R. Dokmeci, Y. Yang, A. Khademhosseini, Lab. Chip 2014, 14, 2202.

[39] K. Yue, G. Trujillo-de Santiago, M. M. Alvarez, A. Tamayol, N. Annabi, A. Khademhosseini, Biomaterials 2015, 73, 254.

[40] B. J. Klotz, D. Gawlitta, A. J. W. P. Rosenberg, J. Malda, F. P. W. Melchels, Trends Biotechnol. 2016, 34, 394.

[41] E. Meijering, Cytometry A 2010, 77A, 693.

[42] N. M. Radio, W. R. Mundy, NeuroToxicology 2008, 29, 361.

[43] A. Odawara, H. Katoh, N. Matsuda, I. Suzuki, Sci. Rep. 2016, 6, 26181.

[44] A. M. Winans, S. R. Collins, T. Meyer, eLife 2016, 5, e12387.

[45] H. J. Lim, Z. Khan, T. S. Wilems, X. Lu, T. H. Perera, Y. E. Kurosu, K. T. Ravivarapu, M. C. Mosley, L. A. Smith Callahan, ACS Biomater. Sci. Eng. 2017, 3, 776.

[46] J. F. Cherry, N. K. Bennett, M. Schachner, P. V. Moghe, Acta Biomater. 2014, 10, 4113.

[47] H. J. Lim, Z. Khan, T. S. Wilems, X. Lu, T. H. Perera, Y. E. Kurosu, K. T. Ravivarapu, M. C. Mosley, L. A. Smith Callahan, ACS Biomater Sci Eng 2017, 3, 776.

[48] K. J. Tomaselli, K. M. Neugebauer, J. L. Bixby, J. Lilien, L. F. Reichard, Neuron 1988, 1, 33.

[49] H. Togashi, K. Abe, A. Mizoguchi, K. Takaoka, O. Chisaka, M. Takeichi, Neuron 2002, 35, 77.

[50] Z. J. Tan, Y. Peng, H. L. Song, J. J. Zheng, X. Yu, Proc Natl Acad Sci 2010, 107, 9873.

[51] B. Du, C. Zeng, W. Zhang, D. Quan, E. Ling, Y. Zeng, J. Biomed. Mater. Res. A 2014, 102, 1715.

[52] G. Li, M.-T. Che, K. Zhang, L.-N. Qin, Y.-T. Zhang, R.-Q. Chen, L.-M. Rong, S. Liu, Y. Ding, H.-Y. Shen, S.-M. Long, J.-L. Wu, E.-A. Ling, Y.-S. Zeng, Biomaterials 2016, 83, 233.

[53] S. Liu, X. Sun, T. Wang, S. Chen, C. Zeng, G. Xie, Q. Zhu, X. Liu, D. Quan, Mater. Sci. Eng. C 2018, 83, 130.

[54] T. Ren, B. Grosshäuser, K. Sridhar, T. J. F. Nieland, A. Tocchio, U. Schepers, U. Demirci, Biomaterials 2019, 197, 171.

