## Supplemental Figures for "Development of an N-Cadherin Biofunctionalized Hydrogel to Support the Formation of Synaptically Connected Neural Networks"

Inventory

Supplemental Figure 1

Supplemental Figure 2

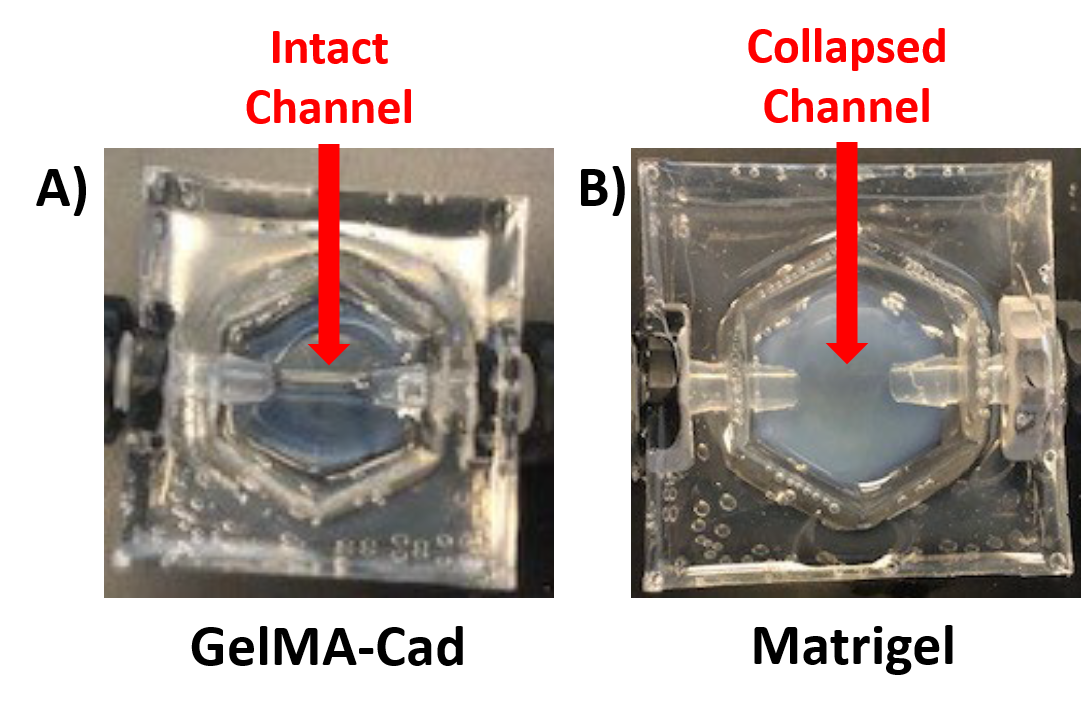

**Supplemental Figure 1:** **Assessment of patterned architectures in hydrogels fabricated from GelMA-Cad or Matrigel.** PDMS molds were filled with GelMA-Cad or Matrigel and crosslinked around a piece of silicone tubing, which was then manually removed. **(A)** GelMA-Cad hydrogel shows an intact channel that can be perfused. **(B)** The channel in the Matrigel hydrogel collapses after the tubing is removed.

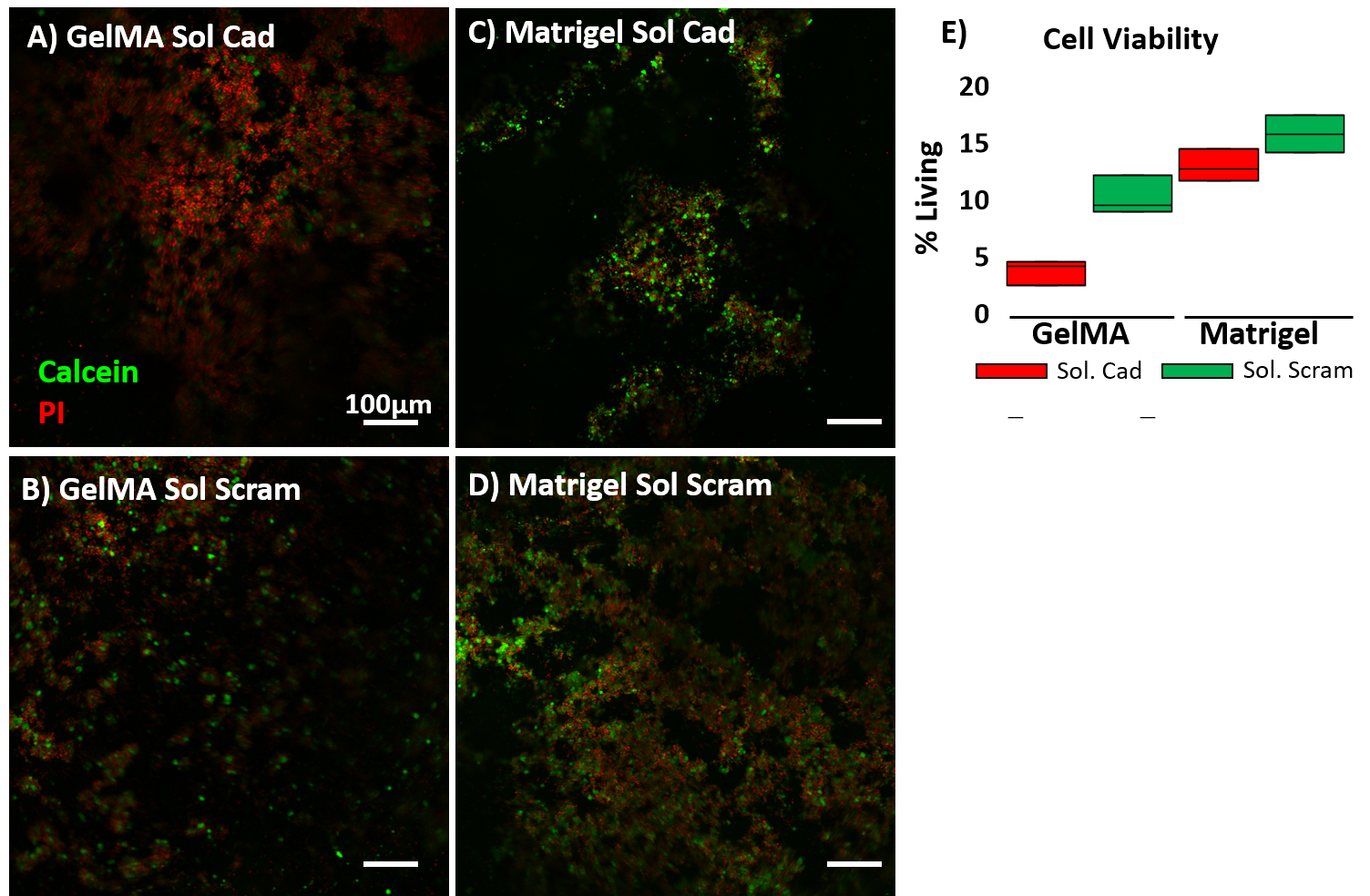

**Supplemental Figure 2:** **Assessment of cell viability in iPSC-derived neurons embedded in GelMA or Matrigel with soluble peptides.** For panels A-D, cells were labeled with calcein (green) to visualize live cells and propidium iodide (PI, red) to visualize dead cells. All images were taken 4 days after embedding. **(A)** Neurons embedded in GelMA with soluble N-cadherin peptide. **(B)** Neurons embedded in GelMA with soluble scrambled peptide. **(C)** Neurons embedded in Matrigel with soluble N-cadherin peptide. **(D)** Neurons embedded in Matrigel with soluble scrambled peptide. **(E)** Quantification of cell viability. Data represent mean ± S.D. from 3 biological replicates, with 4 images assessed per replicate.
